# Structure of the ALS Mutation Target Annexin A11 Reveals a Stabilising N-Terminal Segment

**DOI:** 10.1101/2020.03.27.011445

**Authors:** Peder August Gudmundsen Lillebostad, Arne Raasakka, Silje Johannessen Hjellbrekke, Sudarshan Patil, Trude Røstbø, Hanne Hollås, Siri Aastedatter Sakya, Peter D. Szigetvari, Anni Vedeler, Petri Kursula

**Author notes:** These authors contributed equally (Peder August Gudmundsen Lillebostad, Arne Raasakka).

## Abstract

The functions of the annexin family of proteins involve binding to Ca^2+^, lipid membranes, other proteins, and RNA, and the annexins share a common folded core structure at the C terminus. Annexin A11 (AnxA11) has a long N-terminal region, which is predicted to be disordered, binds RNA, and forms membraneless organelles involved in neuronal transport. Mutations in AnxA11 have been linked to amyotrophic lateral sclerosis (ALS). We studied the structure and stability of AnxA11 and identified a short stabilising segment in the N-terminal end of the folded core, which links domains I and IV. Crystal structure of the AnxA11 core highlights main-chain hydrogen bonding interactions formed through this bridging segment, which are likely conserved in most annexins. The structure was also used to study the currently known ALS mutations in AnxA11. Three of these mutations correspond to buried Arg residues highly conserved in the annexin family, indicating central roles in annexin folding. The structural data provide starting points for detailed structure-function studies of both full-length AnxA11 and the disease variants being identified in ALS.

## 1. Introduction

The annexin (Anx) family of Ca^2+^-binding proteins is widely distributed in eukaryotes, with twelve Anxs found in vertebrates [1–3]. All Anxs share multiple structural elements, most notably an evolutionarily conserved C-terminal Anx core. This core consists of four similar domains, each comprised of ~70 amino acids arranged into five *α* helices, termed A-E [1,4]. The only exception is AnxA6, which contains eight domains, most likely due to a fusion of duplicated *anx*A5 and *anx*A10 genes [5]. The five helices within each domain are connected *via* short loops, forming a right-handed superhelix. Helices A and B as well as helices D and E are oriented pairwise in an antiparallel fashion, while helix C is positioned as a bridge orthogonally to the others [4]. The loops connecting helix A to helix B and helix D to helix E contain the Ca^2+^ binding sites [1,6].

The Anxs are structurally distinct from the EF-hand and C2-domain Ca^2+^-binding proteins [2]. The Anxs reversibly bind negatively charged lipids in a Ca^2+^-dependent manner, which is a defining feature of this protein family [1]. The binding is mediated through the convex side of the core structure, coupled to a conformational change induced by Ca^2+^ binding [1,2]. Through lipid binding, Anxs can affect important proteolipid membrane properties, such as curvature and phase separation [7,8].

While the folded C-terminal core structure of the vertebrate Anxs is conserved, the structure of the N terminus varies greatly between different Anxs and determines their individual properties [1]. This region typically contains 10 - 30 residues. However, AnxA7 and AnxA11 harbour unusually long N termini. AnxA11 contains the longest N-terminal domain of all Anxs, with ~200 residues [9]. The N termini of AnxA7 and AnxA11 resemble each other, both being hydrophobic, enriched in Gly, Tyr, and Pro [10–12]. The Ca^2+^-dependent interaction of AnxA11 with phosphatidylserine changes the conformation of its N terminus [13]. Furthermore, the AnxA11 N terminus interacts with the apoptosis-linked gene-2 protein (ALG-2) [14] and S100A6 (calcyclin) [15].

Rabbit AnxA11 appears to exist as two isoforms; AnxA11A and AnxA11B [16]. Only the former binds S100A6 in the presence of Ca^2+^ *via* the region Gln49-Thr62. This region is absent in AnxA11B [16–18], suggesting isoform-specific functions. S100A6 also binds to AnxA1 and AnxA2 in addition to AnxA11 [19,20], indicating functional overlap between these three proteins.

In certain cell types, AnxA11 is predominantly localised to the cytoplasm, while in others, it is mainly present in the nucleus [15,21]. It is likely that the A isoform is enriched in the nucleus, whereas the B isoform is specifically localised to the cytoplasm [16]. The N terminus was suggested to be responsible for the nuclear localisation [18,22]. In the nucleus, AnxA11 appears to be involved in events related to the cell cycle. It participates in cytokinesis, playing an essential role in the formation of the midbody. Cells lacking AnxA11 cannot complete cytokinesis and undergo apoptosis [23].

Expression levels of AnxA11 have been linked to various cancers [24–29]. Recent studies have revealed that mutations in AnxA11 play an important role in the development of the neurodegenerative disease amyotrophic lateral sclerosis (ALS) [8,30–36]. The molecular basis of these mutations is unclear, as an experimentally determined 3D structure of AnxA11 has been lacking.

Since the long N terminus of AnxA11 renders recombinant production of the protein challenging, N-terminally truncated recombinant forms were generated, and the crystal structure of one form, Δ188AnxA11, was determined. We show that a 4-residue sequence motif in the N terminus of the core structure, conserved in most Anxs, functions as a Velcro-like bridging segment and is important for the thermal stability of the Anx core structure. The known ALS mutations are analysed based on the crystal structure, providing a view into conserved features of the Anx core fold.

## 2. Materials and Methods

### 2.1. Cloning of full-length and N-terminally truncated forms of AnxA11

In order to obtain full-length rat AnxA11, RT-PCR was performed using total RNA isolated from PC12 cells by the Trizol method [37]. The cDNA of AnxA11 was obtained using the iScript kit (Bio-Rad) according to the manufacturer’s instructions and the forward primer 5’-ATCCGG<u>CCATGG</u>GTATGAGCTATCCAGGCTATCCAC-3’ (*Nco*I restriction site is underlined) and the reverse primer 5’- ATCCGG<u>GGTACC</u>TCAGTCGTTGCCACCACAGATC-3’ (*Acc*65I restriction site is underlined). The *anx*A11 cDNA was digested with *NcoI* and *Acc*65I, purified after separation in a 1% agarose gel, and ligated into the same restriction sites of the pETM10 vector (a kind gift from Gunter Stier, Heidelberg). Two truncated versions of AnxA11 were generated by PCR using the pETM10 plasmid harbouring rat full-length *anx*A11 cDNA as a template and using the same reverse primer as for full-length AnxA11. The forward primers were 5’-ATCCGG<u>CCATGG</u>GTAGAGGCACCATCA-3’ and 5’-ATCCGG<u>CCATGG</u>GTACCGATGCTTCTG - 3’(the *NcoI* restriction site is underlined), resulting in Δ188AnxA11 (starting with 189-RGTI) and Δ192AnxA11 (starting with 193-TDAS), respectively. Both constructs code for the tag sequence MLHHHHHHPMG at the N terminus. The cDNAs were ligated into the pETM10 vector after digestion of PCR fragments with *Acc*65I and *NcoI*. Constructs were verified by DNA sequencing.

### 2.2. Expression and purification of full-length and N-terminally truncated forms of AnxA11

*E. coli* BL21(DE3) were transformed with AnxA11 expression plasmids. The bacteria were grown in 5 ml LB medium with 50 μg/ml kanamycin until an OD_600_ of 0.6, after which 1 mM isopropyl β-D-1-thiogalactopyranoside (IPTG) was added to induce protein expression at +25 °C for 4 h or at +15 °C overnight (ON), to determine optimal expression conditions. At the end of protein expression, 1 ml of the culture was withdrawn and centrifuged at 2000 g for 10 min at +4 °C. The supernatant was discarded, and the pellet was resuspended in 500 μl lysis buffer (50 mM Na_2_HPO_4_, 500 mM NaCl, 5% (v/v) glycerol, 10 mM imidazole) containing 0.5 μg/ml DNase I (Sigma), 0.25 μg/ml RNase A (Sigma), and EDTA-free protease inhibitor cocktail (cOmplete, Roche). The samples were sonicated on ice, whereafter 100-μl aliquots (total lysate) were withdrawn. Additional 100-μl aliquots of the lysates were transferred into new tubes and centrifuged at 21000 *g* for 15 min at +4 °C. The supernatant was withdrawn. The pellet was resuspended in 100 μl lysis buffer. After addition of denaturation buffer, 20 μl of the protein samples were separated on 4-20% SDS-PAGE (Mini-PROTEAN TGX Precast Protein Gels, Bio-Rad) at 20 mA and 250 V as the limiting voltage.

For large-scale purification, the above procedure was repeated using 300 ml LB medium. The full-length (FL) and N-terminally truncated AnxA11 were all induced at +25 °C for 4 h, while FL-AnxA11 was also expressed ON at +15 °C. The cells were collected by centrifugation at 4000 *g* for 15 min. The pellets were frozen at −80 °C in lysis buffer before the addition of 0.5 μg/ml DNase I, 0.25 μg/ml RNase A, and EDTA-free protease inhibitor cocktail (cOmplete, Roche). Subsequently, cells were broken by sonication, and the lysates were centrifuged at 16000 g for 30 min at +4 °C. All purification steps were performed at +4 °C.

We took advantage of an N-terminal His-tag to purify AnxA11. In addition, a His-tag protects the protein from degradation at the N terminus [13]. Purification on Co^2+^ instead of Ni^2+^ resin resulted in higher purity. The lysate supernatant was loaded onto a Co^2+^-NTA agarose column and incubated for 30 min. Subsequently, the proteins on the Co^2+^ resin were washed with equilibration buffer (50 mM Na_2_HPO_4_, 0.3 M NaCl, 10 mM imidazole; pH 8) and high-salt buffer (50 mM Na_2_HPO_4_, 1 M NaCl, 10 mM imidazole; pH 8), both buffers containing EDTA-free protease inhibitors (cOmplete, Roche). Elution buffer (50 mM Na_2_HPO_4_, 0.3 M NaCl, 250 mM imidazole; pH 8) containing EDTA-free protease inhibitors (cOmplete, Roche) was used to elute the proteins. After adding EGTA to 2 mM, the eluates were quickly transferred onto PD-10 columns for a buffer exchange to 20 mM Tris, pH 8. All recombinant forms of AnxA11 were subjected to size exclusion chromatography using a Superdex 75 or 200 Increase 10/300 GL column (GE Healthcare). Protein concentration was determined by absorbance at 280 nm (using molecular masses of 55513, 36828, and 36401 Da as well as sequence-based extinction coefficients of 42750, 13410, and 13410 M^−1^ cm^−1^ for full-length AnxA11, Δ188AnxA11, and Δ192AnxA11, respectively). The protein size and purity were confirmed by SDS-PAGE and Coomassie Brilliant Blue staining.

### 2.3. Circular dichroism spectroscopy

A Jasco J-810 spectropolarimeter with a Peltier temperature control unit was used for far-UV circular dichroism (CD) spectrocopy. Melting curves were recorded from +25 to +75 °C at 222 nm, at a heating rate of 40 °C/h, to determine the thermal transition temperature (T_m_). T_m_ was estimated in GraphPad Prism using four-parameter logistic regression.

Synchrotron radiation CD (SRCD) data were acquired from 0.5 mg/ml Δ188AnxA11 and Δ192AnxA11 in 20 mM Tris-HCl, pH 8.0 on the AU-CD synchrotron beamline at ASTRID2 (ISA, Aarhus, Denmark). Samples containing 1 mM CaCl_2_ were included in addition to CaCl_2_-free samples to detect Ca^2+^-induced changes in the secondary structure content. Closed 100-μm circular cells were used (Suprasil, Hellma Analytics). Spectra were recorded from 170 to 280 nm at +10 °C. Baselines were subtracted, and CD units were converted to Δε (M^−1^ cm^−1^) in CDtoolX [38]. The spectra were truncated based on the HT voltage and the CD spectrum noise levels.

### 2.4. Small-angle X-ray scattering

SAXS data were collected from 0.41 – 2.96 mg/ml Δ188AnxA11 and Δ192AnxA11 in 20 mM Tris-HCl on the EMBL/DESY P12 beamline, PETRA III (Hamburg, Germany) [39]. Fixed protein concentration series at 0.7 – 1.0 mg/ml were measured with 0 – 1 mM CaCl_2_ to follow potential oligomerisation. Monomeric bovine serum albumin (66.5 kDa) was used as a molecular weight standard. Data were processed and analysed using the ATSAS package [40], and GNOM was used to calculate distance distribution functions [41]. Theoretical scattering of the crystal structure was calculated and fitted against the SAXS data using CRYSOL [42]. *Ab initio* modelling was performed using GASBOR [43]. SUPCOMB [44] was used for crystal structure and *ab initio* model superposition. SAXS data collection, processing, and fitting parameters are presented in Table 1.

**Table 1.** Small-angle X-ray scattering parameters and analysis.

| Data collection parameters |  |  |
| --- | --- | --- |
| Instrument | P12, PETRAIII, DESY |  |
| Wavelength (nm) | 0.124 |  |
| Angular range (nm<sup>-1</sup>) | 0.022 - 7.33 |  |
| Exposure time (s) | 0.045 |  |
| Exposure temperature (°C) | 10 |  |
| Protein | AnxA11Δ188 | AnxA11Δ192 |
| Concentration range (mg/ml) | 0.77 - 2.96 | 0.41 - 1.18 |
| Structural parameters |  |  |
| *I*<sub>0</sub> (relative) [from Guinier] | 0.02759 | 0.0274 |
| *R*<sub>g</sub> (nm) [from Guinier] | 2.44 | 2.43 |
| *I*<sub>0</sub> (relative) [from P(r)] | 0.02692 | 0.02748 |
| *R*<sub>g</sub> (nm) [from P(r)] | 2.500 | 2.503 |
| *D*<sub>max</sub> (nm) [from GNOM] | 10.00 | 9.97 |
| *V*<sub>Porod</sub> (nm<sup>3</sup>) [from GNOM] | 58.03 | 59.53 |
| Molecular mass determination |  |  |
| Molecular mass *M*<sub>r</sub> (kDa) [from *I*<sub>0</sub> using Guinier]* | 34.3 | 34.1 |
| Molecular mass *M*<sub>r</sub> (kDa) [from *I*<sub>0</sub> using P(r)]<sup>1</sup> | 33.5 | 34.2 |
| Molecular mass *M*<sub>r</sub> (kDa) [from *V*<sub>Porod</sub>] | 34.1 | 35.0 |
| Molecular mass *M*<sub>r</sub> (kDa) [from absolute scale] | 34.8 | 34.8 |
| Theoretical *M*<sub>r</sub> from sequence (kDa) | 36.8 | 36.4 |
| Software |  |  |
| Primary data reduction & processing | PRIMUS |  |
| *Ab initio* modelling | GASBOR |  |
| χ<sup>2</sup> | 0.770 | 0.629 |
| Fitting of theoretical scattering curve to data | CRY SOL |  |
| χ<sup>2</sup> | 0.888 | 0.749 |

<sup>1</sup> Monomeric BSA standard was used as reference (*M*<sub>r</sub> = 66.5 kDa; *I*<sub>0</sub> = 0.0534)

### 2.5. Protein crystallisation, data collection, and structure determination

Sitting-drop vapour diffusion crystallisation experiments were set up in 96-well format. X-ray diffraction data were collected on the EMBL P13 beamline, PETRA III (Hamburg, Germany) [45] from a crystal grown in 100 mM Bis-tris (pH 5.5), 3 M NaCl at +20 °C; the drop contained 150 nl of 12.7 mg/ml Δ188AnxA11 and 150 nl of reservoir solution. The crystal was cryoprotected by adding a solution containing 25% glycerol and 75% reservoir solution directly onto the crystallization drop. Upon mounting, the crystal was flash-frozen using liquid N_2_, and kept at 100 K during X-ray diffraction data collection.

Diffraction data were processed with XDS [46], and the structure was solved by molecular replacement in Phaser [47] with PDB entry 1ANN (bovine AnxA4) as search model [48]. Refinement was carried out using phenix.refine [49] and model building with coot [50]. The structure quality was assessed with MolProbity [51]. The data processing and refinement statistics are presented in Table 2.

**Table 2.** Crystallographic data collection and refinement.

| Construct | $\Delta 188\text{AnxA11}$ |
| --- | --- |
| Wavelength (Å) | 0.9763 |
| Space group | P1 |
| Unit cell parameters | $a = 39.1 \text{ Å}, b = 86.7 \text{ Å}, c = 87.3 \text{ Å}$<br>$\alpha = 114.0^\circ, \beta = 102.0^\circ, \gamma = 97.2^\circ$ |
| Resolution range (Å) | 50 – 2.30 (2.36 – 2.30)<sup>1</sup> |
| Completeness (%) | 95.4 (95.0) |
| $\langle I/\sigma I \rangle$ | 5.2 (1.1) |
| $R_{\text{sym}}$ (%) | 17.5 (164.0) |
| $R_{\text{meas}}$ (%) | 20.9 (196.6) |
| $CC_{1/2}$ (%) | 99.2 (47.6) |
| redundancy | 3.3 (3.2) |
| $R_{\text{cryst}}/R_{\text{free}}$ (%) | 22.8 (26.6) |
| Rmsd bond length (Å) | 0.002 |
| Rmsd bond angle (°) | 0.4 |
| MolProbity score (percentile) | 1.58 (98<sup>th</sup>) |
| Ramachandran favoured/disallowed (%) | 97.2/0.3 |

<sup>1</sup> Numbers in parentheses refer to the highest-resolution shell.

For structure analysis, the programs PyMOL [52] and APBS [53] were used. The AnxA11 crystal structure was deposited at the PDB with the entry code 6TU2.

## 3. Results and Discussion

### 3.1. Solubility assays, large scale expression and purification of the N-terminally truncated forms of AnxA11

Induction was performed at +25 °C for 4 h and at +15 °C ON to determine optimal expression conditions. There was little or no difference in yield between the two modes of expression. FL-AnxA11 was consistently more challenging to express than its truncated forms, yielded remarkably low concentrations and was notoriously prone to aggregation, as reported previously by others [13]. The solubility of the truncated forms of AnxA11 in bacterial lysates was assessed after centrifugation at 21000 *g* by comparing samples from the total bacterial lysate with the corresponding supernatant and pellet by SDS-PAGE (Figure 1). In contrast to FL-AnxA11, both Δ188AnxA11 and Δ192AnxA11 were mainly soluble (Figure 1A). FL-AnxA11 as well as the N-terminally truncated Δ188AnxA11 and Δ192AnxA11 were purified by affinity chromatography on Co^2+^-resin, which greatly improved the purity of the recombinant both proteins compared to Ni^2+^-based chromatography. The N-terminal truncations resulted in a marked enhancement in expression, giving a five-to-tenfold increase in yield compared to FL-AnxA11 (Figure 1B). This suggests that the N terminus is responsible for the aggregation of the full-length protein. This may be related to the observed phase separation phenomena linked to the N-terminal segment [8], or misfolding of the protein in bacterial cells.

**Figure 1.**
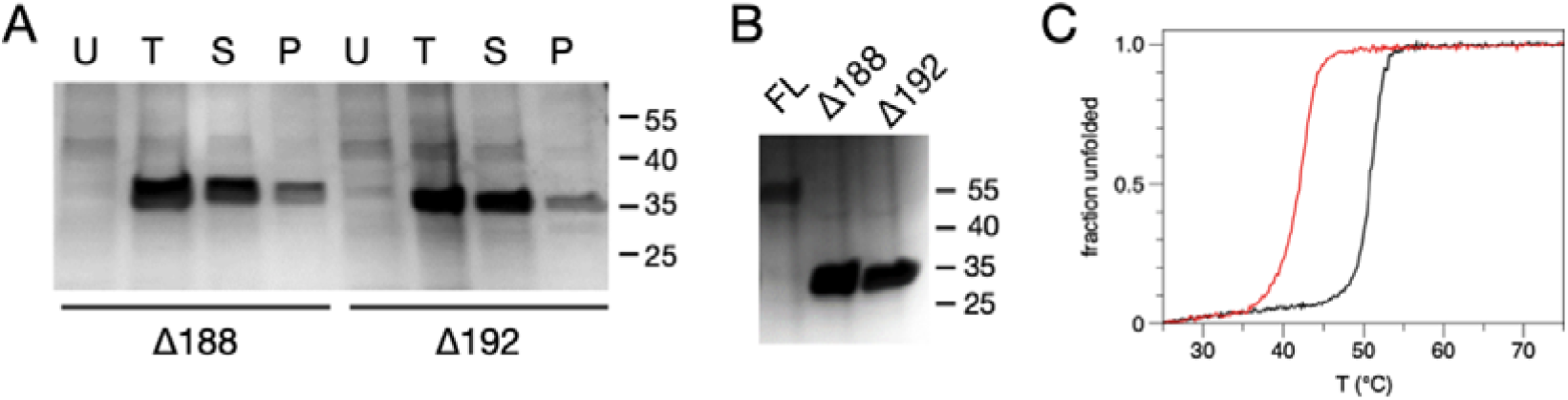
The solubility and purification of N-terminally truncated forms of AnxA11 expressed at 15 °C overnight and their thermal stability. (A) Uninduced (U), total (T), supernatant (S), and pellet (P) fractions of bacterial lysates containing Δ188AnxA11 or Δ192AnxA11 as indicated. (B) 10 μl of FL-AnxA11 and the N-terminally truncated forms Δ188AnxA11 and Δ192AnxA11, purified using Co^2+^-resin. Molecular mass standards are indicated to the right. Proteins were separated in 4-20% SDS gradient gels and Coomassie-stained. (C) CD-monitored thermal disruption of α-helicity of 20 μM Δ188AnxA11 (black) and Δ192AnxA11 (red).

### 3.2 Thermal stability of the N-terminally truncated Δ188AnxA11 and Δ192AnxA11

To address thermal stability, ellipticity at 222 nm was monitored as a function of temperature for Δ188AnxA11 and Δ192AnxA11. The thermal transition temperatures were estimated from the inflection points of the melting curves (Figure 1C). The T_m_ of FL-AnxA11 was earlier reported to be +45 °C [13]. The T_m_ of Δ188AnxA11 showed a T_m_ of +50.76 ± 0.02°C, while that of Δ192AnxA11 was substantially lower (+41.87± 0.02°C). This difference highlights the importance of the four amino acid residues 189-RGTI-192 in thermally stabilising the core structure of AnxA11. Cooperative unfolding of the two truncated forms of AnxA11 remained largely unchanged but decreased slightly in Δ192AnxA11. This increased thermal stability of Δ188AnxA11 compared to Δ192AnxA11 may explain why we obtained crystals of Δ188AnxA11, but not of the other N-terminally truncated form. The dramatic difference in T_m_ is necessarily accounted for by at least one of the four additional residues present in Δ188AnxA11 being involved in the stabilising of the core structure.

### 3.3. The crystal structure of AnxA11

Using Δ188AnxA11, we solved the crystal structure of the rat AnxA11 core domain. The asymmetric unit contains 3 molecules of AnxA11, and the structure presents the canonical annexin core fold. The Ca^2+^ binding sites are partially occupied, whereby 2 monomers have 3 Ca^2+^ ions bound, and one monomer has 4 (Figure 2). As is typical for the Anxs, all Ca^2+^ binding sites are on one face of the molecule, promoting calcium-dependent lipid membrane binding (Figure 2).

**Figure 2.**
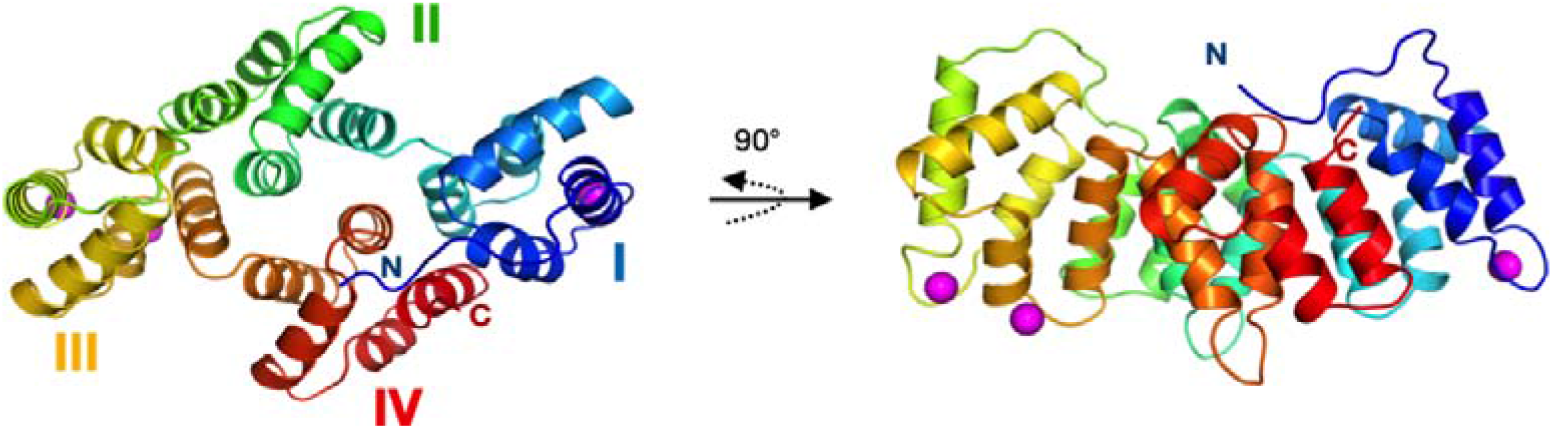
Crystal structure of the AnxA11 core domain. The 4 subdomains are numbered and coloured from blue to red. The N and C termini are indicated, and bound Ca^2+^ ions are shown in magenta. Shown is one monomer of the 3 in the asymmetric unit.

The substantial difference in thermal stability induced by the 4 N-terminal residues of the Δ188AnxA11construct implies an important structural element at this position. This segment ~10 residues before the AnxA11 domain I helix A are central in the folded structure, making interactions with the termini of several helices of the Anx fold, and effectively enabling a tight contact between the N terminus of domain I and the C terminus of domain IV (Figure 3A,B).

**Figure 3.**
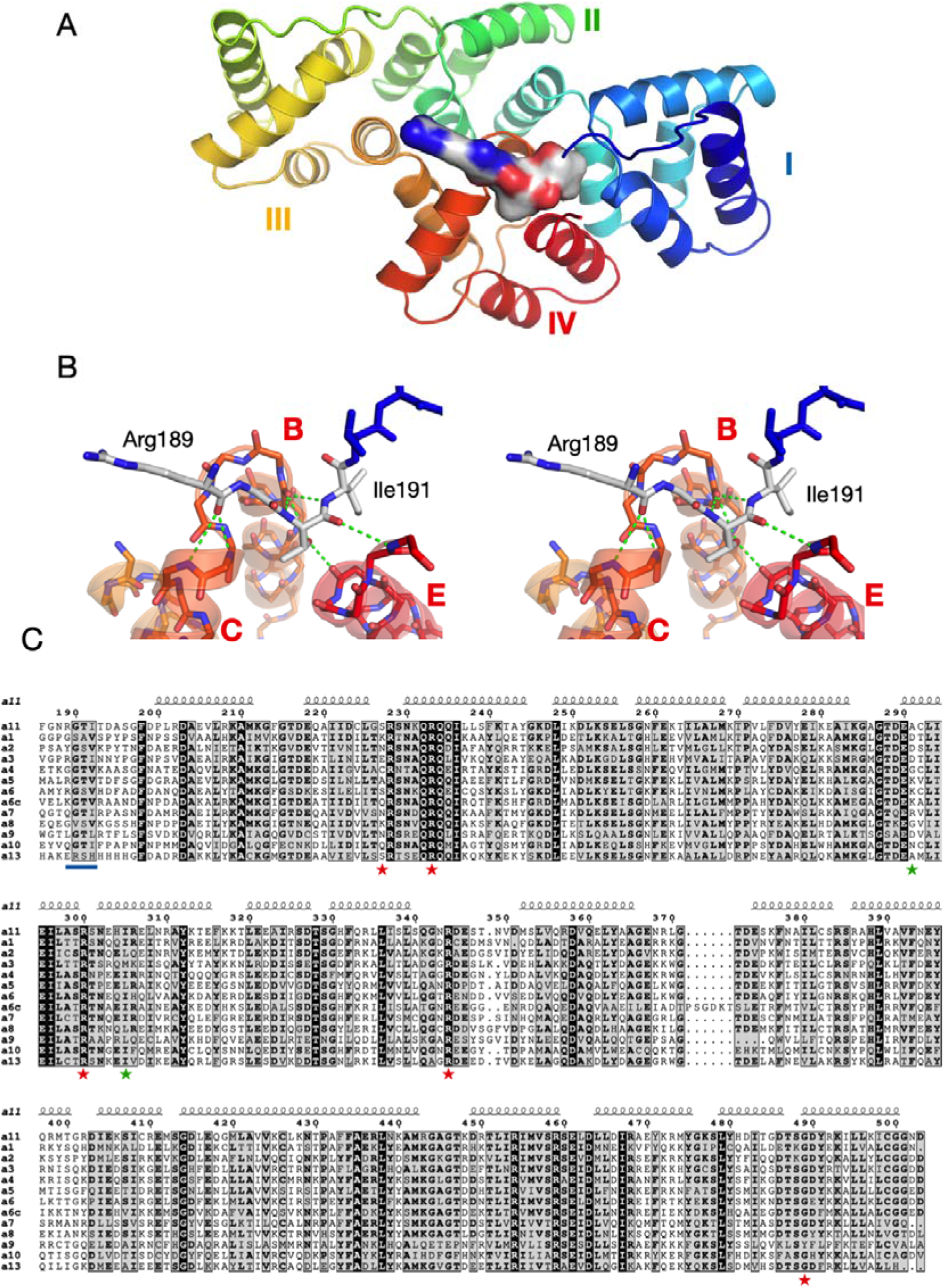
Rat annexins and conservation of the RGTI motif. (A) The N-terminal Velcro-like bridging segment is shown in surface mode, bound to domain IV. (B) Stereo view of the hydrogen bonding interactions of the motif with three helices in domain IV. (C) Sequence alignment of all rat Anxs. The N-terminal motif is underlined in blue, and the mutations (red) and variants (green) linked to ALS are shown with asterisks. A6 and A6c correspond to the N- and C-terminal halves of AnxA6.

Comparisons with other Anxs show that the segment 189-192 is a conserved motif in the Anx family. The only Anx that does not contain the motif at the sequence level is AnxA13; conservation is weak for AnxA1 and AnxA8. The 4-residue segment forms conserved interactions with the ends of helices in domain IV (Figure 3B). These interactions are mainly comprised of main-chain hydrogen bonds, which explains the mode of conservation and the importance of the Gly residue in the motif. Interestingly, it has been proposed that AnxA11 is the common ancestor of all mammalian Anxs [54], except for AnxA7 and AnxA13 – the two phylogenetically oldest Anxs [55]. Thus, the AnxA11 core domain has the highest similarity to the other mammalian Anxs, acting as a sort of prototype [56].

Interestingly, the Tyr23 and Ser25 phosphorylation sites of AnxA2 lie within the same 4-residue region (Figure 3C). Phosphorylation at Tyr23 and Ser25 results in a closed or open conformation, respectively, of AnxA2 [57]. These findings can be understood based on the conserved structure of the observed bridging segment.

AnxA5 has been the most studied Anx on membrane surfaces, on which it forms a crystal-like lattice of trimers [58–61]. The crystal structure of AnxA11 presents trimers very similar to those formed by AnxA5, and hence, could be expected to interact with membranes in a similar manner. In the crystal, the trimers are arranged in layers, with similar contacts as observed for AnxA4 and AnxA5 between the trimers. Figure 4 shows the trimer arrangement of AnxA11; the arrangement in the crystal resembles that of AnxA5 on membranes [58–61]. Like other Anx assemblies, the Ca^2+^ binding sites of the AnxA11 trimer are all on the same face, which would be in direct contact with the membrane (Figure 4A). The membrane-facing surface in the dimer gains a positive charge potential upon Ca^2+^ binding, while the opposite side is relatively featureless (Figure 4B). This observation further highlights the common function of the Anx core structure in binding membrane surfaces, while the N-terminal domain is responsible for most functional differences between the Anxs [1]. One might assume that AnxA11 would arrange on a membrane much like it does in the crystal; the N-terminal segments would then point away from the membrane. This is in line with the observations that AnxA11 induces membrane curvature [7], as well as the suggestion that the N terminus of AnxA11 binds to RNA, while the core structure binds to lipids in lysosomes for long-distance co-transport in neurons [8]. The RNA binding site in AnxA2 has been shown to reside within domain IV [62]. The binding dynamics were recently investigated; AnxA2 binds strongly to an RNA 8-mer (5’-GGGGAUUG-3’) with a K_d_ of 30 nM [63]. The RNA binding site(s) in AnxA11 have not been determined.

**Figure 4.**
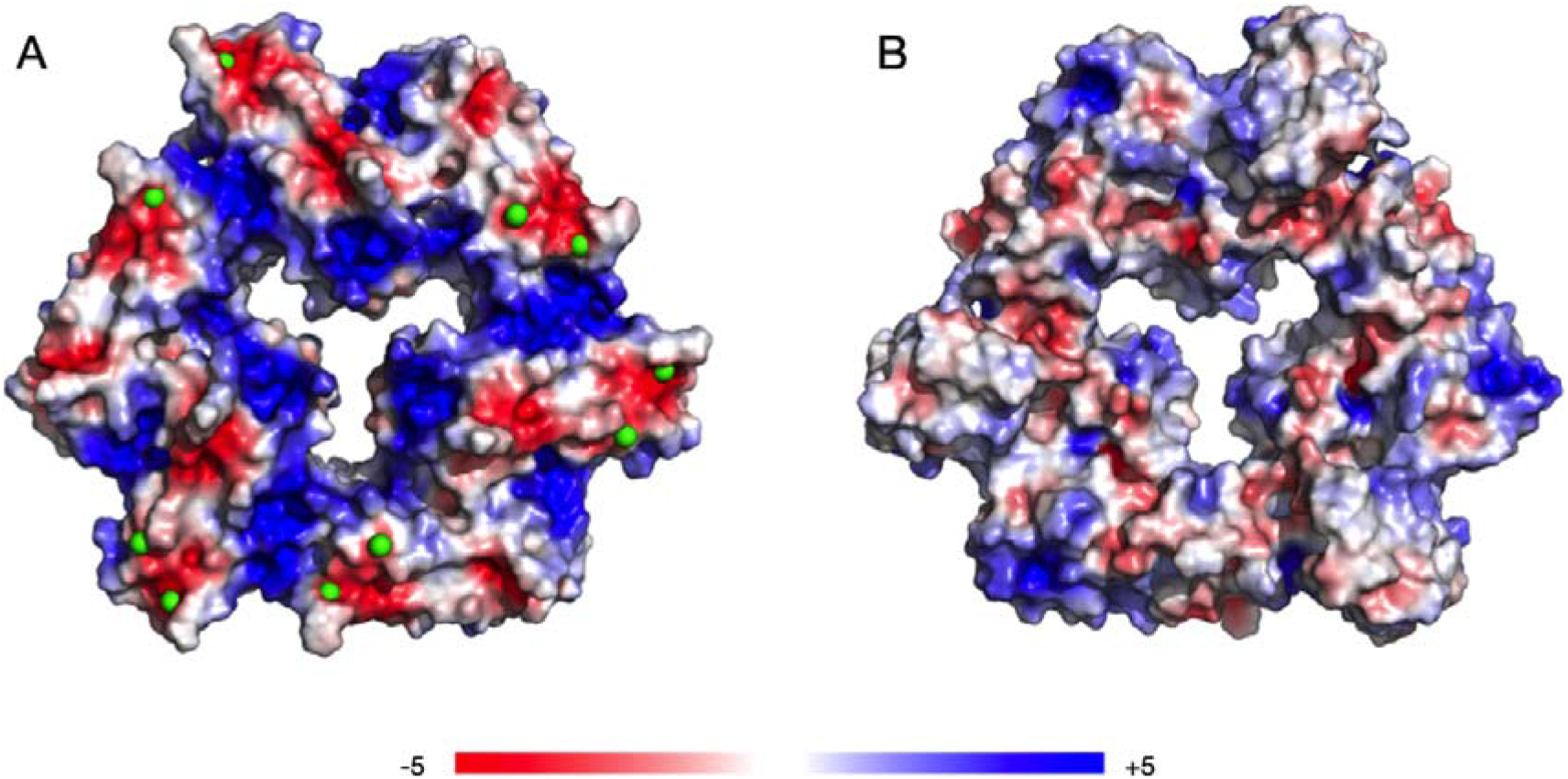
Trimeric assembly of AnxA11 in the crystal, shown as a surface with electrostatic potential. (**A**) View from the Ca^2+^-binding face. Ca^2+^ ions are shown as green spheres. The negative sites are the locations for Ca^2+^ binding, and their neutralization will make the surface very positive, enabling lipid membrane binding. (**B**) The opposite face of AnxA11 is relatively featureless.

### 3.4. Structure-based analysis of ALS disease mutations (ALS)

The crystal structure of rat AnxA11 allows to analyse the structural basis of AnxA11 mutations linked to amyotrophic lateral sclerosis (ALS) [34] in humans. The sequence identity between full-length rat and human AnxA11 is 93.3%, being 96.5% for the core structure. The identification of AnxA11 as an ALS gene is rather recent, and mutations are being continuously identified. Thus, it is safe to assume that more ALS mutations will be identified in the future. We studied the mutations thus far linked to ALS with respect to potential structural implications. Many of the mutations, including the best-characterised D40G [34], lie in the N-terminal extension, which is not present in the crystallised construct. The mutations in this region could affect protein-protein interactions, protein localisation, phase separation, or binding to mRNA [8]. More interesting for the current structural study, however, are the mutations identified within the Anx core domain.

Five mutations have thus far been described in ALS patients within the AnxA11 core: S229R, R235Q, R302C, R346C, and G491R; in addition, G189E is located two residues before the start of the crystallised rat AnxA11 construct [34,36]. In the 3D structure, the AnxA11 mutations can be observed clustered at two locations (Figure 5A). Interestingly, three of these mutations involve a buried Arg residue, and a close look at the alignment reveals that these Arg residues are fully conserved in all rat Anxs. The remaining two mutations replace an exposed, non-conserved, small residue with an Arg. Details of the environment of each of the three conserved Arg residues are shown in Figure 5B-D. In brief, two of them coordinate carbonyl groups at C termini of helices, and one is involved in a conserved extended buried network of salt bridges between domains II and IV. These locations suggest that the ALS mutations involving these conserved Arg residues will have effects on AnxA11 folding and/or stability.

**Figure 5.**
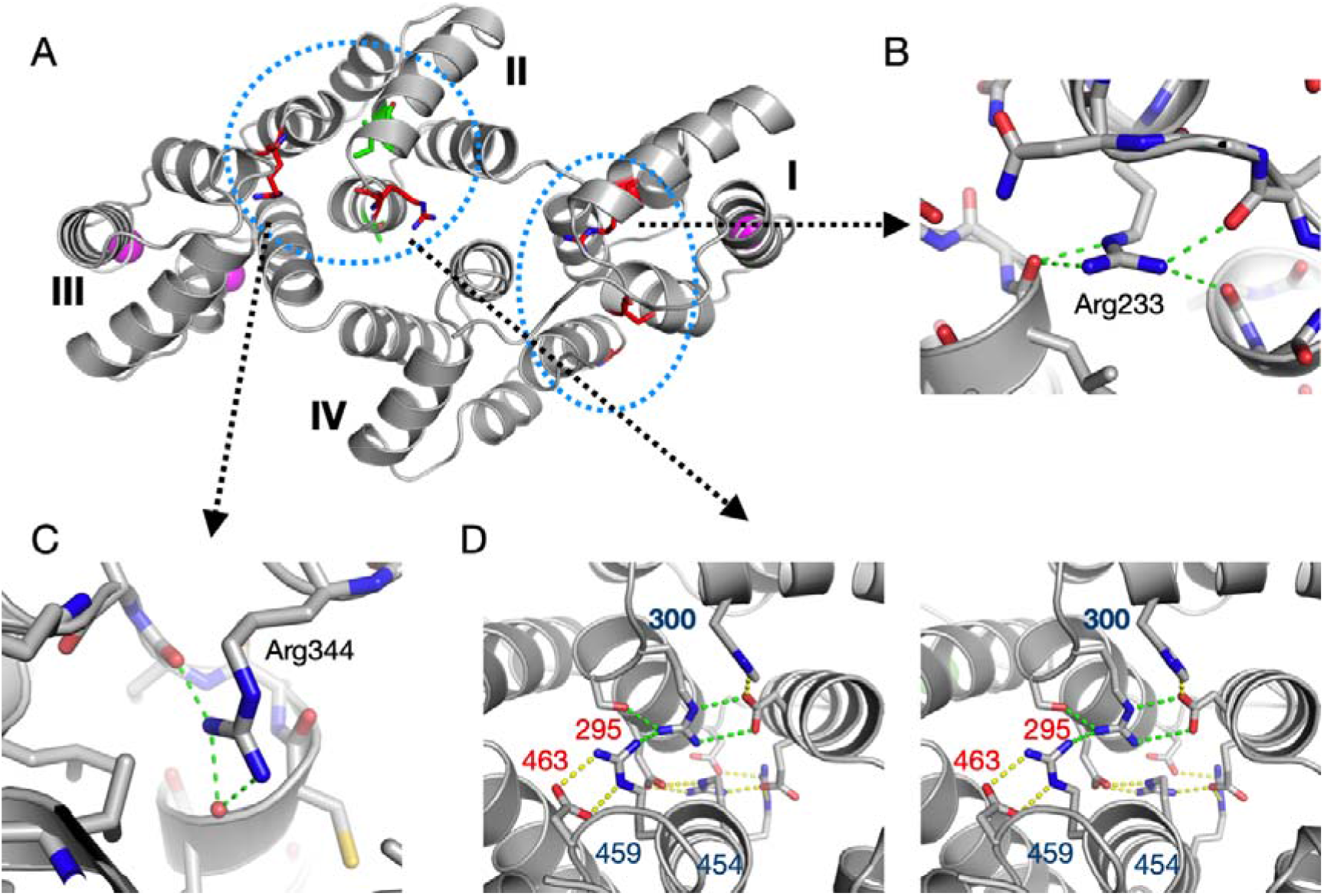
Location of ALS mutations and variants in the AnxA11 structure. (**A**) The five ALS mutations (see text) are highlighted in red and the two rare variants in green. The two 3D clusters of mutations/variants are shown with blue dashed circles. One cluster lies in domain II, possibly affecting folding and/or interactions with domain III. The other cluster is at the interface between domains I and IV, *i.e.* the N and C terminus of the AnxA11 core structure. (**B**) Arg233 (corresponding to the human R235Q mutation) makes 4 contacts to carbonyl groups at the C-terminal end of domain I helices B and E. With its buried positive charge, Arg233 interacts with the negative helix dipole from both helices. (**C**) Arg344 (corresponding to R346C) similarly caps helix B of domain III. (**D**) An extended salt bridge network exists between domains II (top) and IV (bottom) (stereo view). Arg300 (corresponds to R302C) is central in this network. The charged residues with numbers indicated are fully conserved in all rat Anxs (Arg, blue; acidic, red).

ΔΔG predictions for the five mutations on the SDM server [64] indicate decreased stability for all mutations except S229R. Hence, the affected residues may be important for correct folding and/or stability of AnxA11. Recent functional characterisation of the AnxA11 ALS mutation R235Q shows that the mutant protein is prone to aggregation and sequesters FL-AnxA11. This in turn abolishes the binding to calcyclin [34]. In addition, two rare missense variants A293V and I307M were recently discovered in the Chinese Han population [33]. These residues are located in domain II (Figure 5).

The N-terminal tail of AnxA11 has a unique amino acid composition, reminiscent of disordered proteins able to undergo liquid-liquid phase separation (LLPS) [65]. It has the hallmarks of an extended segment poised for *π*-*π* interactions, which are central in LLPS [66]. Indeed, recently the AnxA11 N-terminal region was shown to promote LLPS; its N terminus binds to RNA granules, which do not contain membranes, while the core structure binds to lipids in membranes [8]. Furthermore, ALS-associated mutations in both the N terminus (D40G) and C terminus (R346C) of AnxA11 were shown to promote phase transition from liquid-liquid droplets to more stable gel-like states within AnxA11 droplets and in addition impair the reversal of the states. As a consequence, these mutations disrupt the function of AnxA11 to tether RNA granules with lysosomes [8].

### 3.5. The solution behaviour of the AnxA11 core

The thermal stability difference between Δ188AnxA11and Δ192AnxA11 prompted us to study their conformation in solution using SAXS and SRCD. In SAXS, both variants were monomeric with identical distance distributions and globular folds that superposed well with the crystal structure (Figure 6A-E), indicating that the absence of the RGTI stretch does not lead to large-scale conformational changes. Despite the trimeric packing in the crystal, AnxA11 is monomeric in solution under the employed conditions.

**Figure 6.**
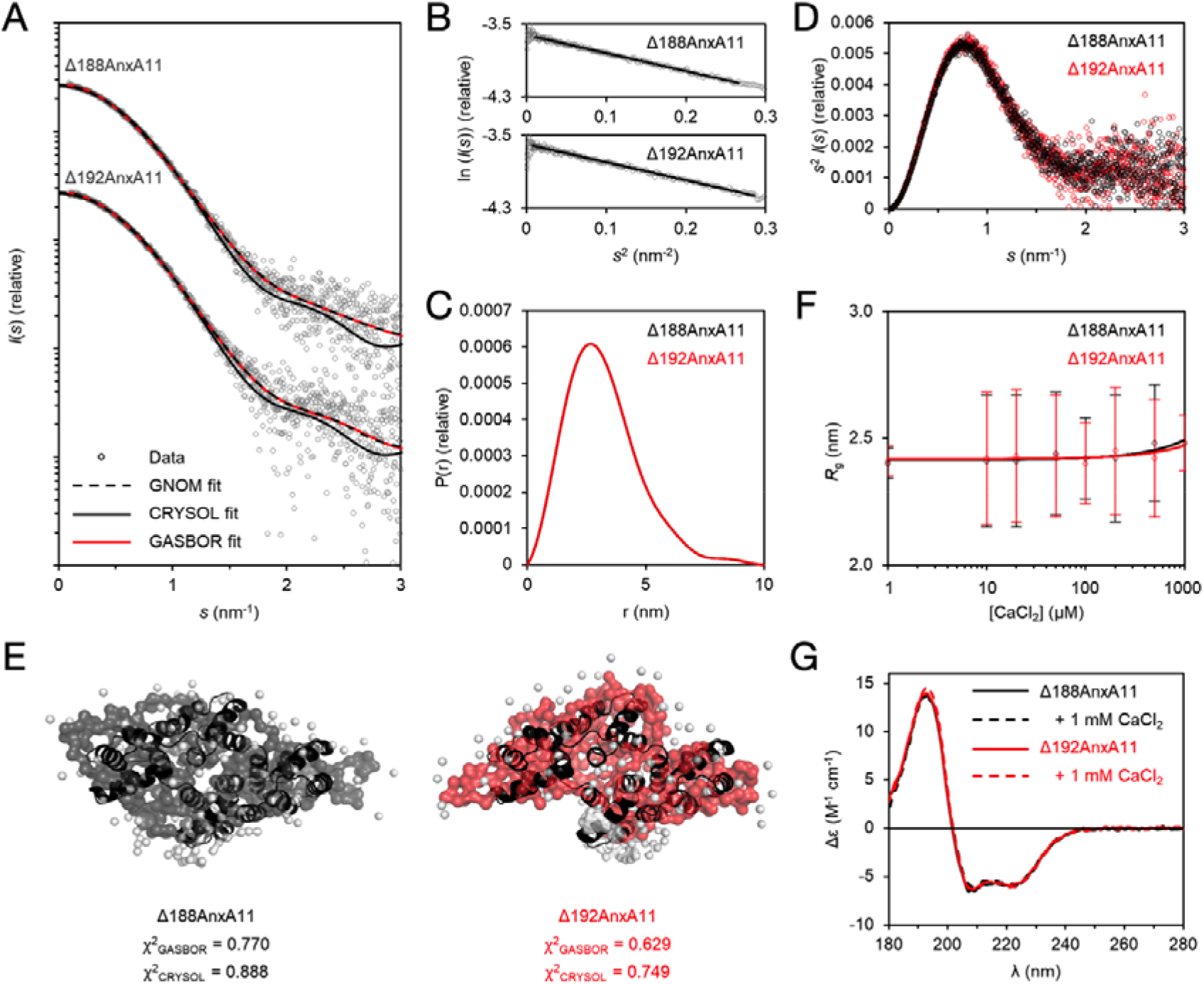
SAXS and SRCD analysis of Δ188AnxA11 and Δ192AnxA11. (**A**) SAXS data for Δ188AnxA11 and Δ192AnxA11 have been offset for clarity. GNOM, GASBOR, and CRYSOL fits are plotted over the measured data to denote the accuracy of distance distribution analysis, *ab initio* modeling, and theoretical scattering calculated from the crystal structure. (**B**) Guinier analysis. (**C**) Distance distribution diagram from GNOM. (**D**) Kratky plot of the SAXS data. (**E**) GASBOR *ab initio* models based on the SAXS data. The crystal structure (black) has been fitted inside. *χ*^2^ values of the GASBOR and CRYSOL fits are indicated. (**F**) R_g_ as a function of CaCl_2_ concentration. The error bars represent the fitting error from Guinier analysis. (**G**) SRCD spectra of Δ188AnxA11 and Δ192AnxA11 in the presence and absence of 1 mM CaCl_2_.

Like most Anxs, AnxA11 has been implicated in Ca^2+^ binding [1,13]. Structural studies of many Anxs have revealed Ca^2+^ binding to the convex side of the core structure, essentially through coordination by acidic residues. To test for effect of Ca^2+^ on the folding of the core structure of AnxA11, we titrated Δ188AnxA11 and Δ192AnxA11 with CaCl_2_ in SAXS and followed the radius of gyration (R_g_; Figure 6F). The R_g_ of both truncated proteins remained the same in up to 1 mM CaCl_2_. Additionally, we carried out SRCD spectroscopy of Δ188AnxA11 and Δ192AnxA11 in the absence and presence of 1 mM CaCl_2_ (Figure 6G). The spectral quality was excellent, with the data being truncated at 178 nm. The presence of Ca^2+^ did not alter the spectra, indicating that Ca^2+^ does not influence the secondary structure content of the core structure. This result confirms that Δ192AnxA11 is folded despite its dramatic decrease in thermal stability, and that the degree of secondary structure content between the two variants is nearly identical. Ca^2+^ induced a large thermal stabilisation of FL-AnxA11, indicated by a rise in T_m_ of ~14 °C [13]. The presence of up to 75 mM Ca^2+^ only induced small increases in *α*-helical content [13]. Thus, Ca^2+^ binding results in only local conformational changes, which primes AnxA11 for lipid membrane binding.

## 4. Conclusions

As the diversity of the vertebrate Anx family is mainly realised through the uniqueness of the N-terminal tail [1,16,67,68], many unique functional features are lost by truncation. However, crucial functional properties of interest, such as mRNA binding, Ca^2+^-dependent binding to phospholipids, as well as Ca^2+^ binding itself, may be retained. Thus, truncated Anxs may serve as a foundation for functional studies, dodging practical issues often encountered due to their long N termini.

The crystal structure of AnxA11, although similar to known Anx core domain structures, provides important starting points to further understand the structure-function relationships in AnxA11 and other Anx family members. The identification of a strongly stabilizing Velcro-like bridging segment at the N terminus of the Anx core, as well as the fact that many known ALS mutations correspond to highly conserved buried Arg residues in the Anx fold, provide new insights into Anx folding. AnxA11, like many other ALS targets, is part of a membraneless organelle, a ribonucleoprotein complex, involved in neuronal transport of mRNA granules [8,69–71]. It is evident that this process can be affected by mutations in either the N-terminal region or the folded core domain of AnxA11, which in turn can affect binding to both RNA, lipid membranes, and Ca^2+^. The fact that the bridging segment is a target for phosphorylation in AnxA2 shows that in addition to simply stabilizing the core structure, this segment may be a conformational regulatory element common to most Anxs.

## Author Contributions

Conceptualization, A.V. and P.K.; methodology, P.A.G.L., A.R., and A.V.; validation, A.R. and P.K.; formal analysis, A.R. and P.K.; investigation, P.A.G.L., A.R., S.J.H., S.P., T.R., H.H., S.A.S., P.D.S., A.V., and P.K.; resources, P.A.G.L., S.J.H., S.P., T.R., H.H., S.A.S., P.D.S., and A.V.; data curation, A.R. and P.K.; writing—original draft preparation, P.A.G.L., A.R., and A.V.; writing—review and editing, A.R., A.V., and P.K.; visualization, A.R. and P.K.; supervision, A.V. and P.K.; project administration, A.V. and P.K.; funding acquisition, A.V. and P.K. All authors have read and agreed to the published version of the manuscript.

## Funding

This research was funded by the University of Bergen. Synchrotron visits were supported by the EU Framework Programme for Research and Innovation HORIZON 2020 iNEXT and CALIPSOplus programmes, grant numbers 653706 and 730872.

### Acknowledgments

We thank Marianne Goris for valuable assistance in the laboratory. Crystallographic and SAXS data were collected on EMBL Hamburg beamlines P13 and P12, respectively, at the PETRA III storage ring (DESY, Hamburg, Germany). We also thank the SRCD beamline support at the ASTRID2 synchrotron facility (Aarhus, Denmark). We thank the BiSS facility at the University of Bergen for providing vital instrumentation for this study.

